# MULTICLUST – Fast Multinomial Clustering of multiallelic genotypes to infer genetic population structure

**DOI:** 10.1101/2025.08.06.668969

**Authors:** Arun Sethuraman, Wei-Chen Chen, Margaret Wanjiku, Karin S. Dorman

## Abstract

Identifying population structure from multilocus genotype data is key to down-stream genetic analyses, including analysis of genome-wide association (GWAS), genetic genealogy and phylogenetics, in the fields of conservation, forensics, evolution and more. While inference of population structure has been dominated by Bayesian methods, maximum likelihood methods have some benefits in terms of reliability, consistency and efficiency. Here we extend the methods of Tang et al. [53] and Alexander et al. [2] to handle multi-allelic (*e.g*., SNP, STR, allozyme) and polyploid loci with missing data to infer genetic admixture proportions and subpopulation allele frequencies. Comparative analyses of our method, `MULTICLUST`, and `STRUCTURE` [42] on both simulated and empirical data indicate comparable, fast, reproducible, and accurate estimates of population admixture proportions and allele frequencies using `MULTICLUST.MULTICLUST` is implemented in the C programming language and is publicly accessible via www.github.com/arunsethuraman/multiclust.

## 1 Introduction

Inference of genetic population structure has become a primary component of population genetics analyses in a variety of fields, including phylogeography, conservation, epidemiology, trait engineering, genetic genealogy, and forensics [13, 18, 20, 32, 45, 47]. The major goals when inferring population structure are to estimate the number, size, membership, and genetic characteristics of subpopulations based on a sample of multilocus genotypes from a population [30]. Several methods have been proposed to perform such inferences, categorized broadly as non-parametric and model-based methods. Non-parametric methods mostly use Principal Components/Coordinates Analysis (PCA) to identify the presence of structure (EIGENSOFT [38]), and extensions can identify subpopulation membership (`AW-Clust` [16], `PCO-MC` [44], `ipPCA` [23], `SHIPS` [4]). Model-based methods assume genetically distinct populations at equilibrium (Hardy-Weinberg equilibrium and maybe linkage equilibrium) mix in some way to generate a sample [50]. They either take Bayesian (`STRUCTURE` [42], `PARTITION` [8], `BAPS` [7], `HWLER` [39], `STRUCTURAMA` [21], `Spectrum` [51], `StructHDP` [49], `DPART` [37]) or maximum likelihood (`FASTRUCT` [6], `PSMIX` [56], `FRAPPE` [53], `ADMIXTURE` [2]) inferential approaches. PCA-based methods have grown in popularity in the last decade because of theoretical advances [34, 38] and their efficiency on large genomic datasets [27], but model-based methods still dominate in terms of interpretability and extensibility [17].

The Bayesian Markov Chain Monte Carlo (MCMC) approach, particularly `STRUCTURE` [42], has been most popular among molecular ecologists, owing to its ease of use and intuitive visualizations of genetic admixture. However, several recent studies have pointed to issues with (1) speed of computation, (2) reproducibility, and (3) accuracy of inferring genetic population structure using `STRUCTURE`, especially in the context of ecological and evolutionary questions (Gilbert [17], Gilbert et al. [18], Lawson et al. [28]). Any MCMC approach must run to convergence, after discarding potentially large periods of burn-in. Convergence is harder to achieve for large datasets, with many individuals, loci, or alleles per locus [15, 26]. Estimation of the number of subpopulations, *K*, is also difficult with `STRUCTURE`, and despite several efforts to estimate *K* during MCMC [8, 21, 22, 39, 49], these samplers often fail to sample *K* efficiently, especially for large *K* [37].Although Bayesian approaches provide richly informative posterior distributions of admixture proportions, assignments of alleles or individuals to clusters, and in some cases, subpopulation allele frequencies, it can be difficult to interpret the output because the alleles or individuals are exchangeable [8, 21, 24].

Unless posterior distributions are required, maximum likelihood (ML) methods are naturally more efficient because they seek only a point estimate of model parameters [2]. Most ML approaches to estimate population structure [6, 50, 53, 56] use the Expectation-Maximization (EM) algorithm [9] to maximize the likelihood. The EM algorithm is guaranteed to approach a local stationary point [57], but is also slow to converge [9, 35]. Alexander et al. [2] proposed instead a block relaxation algorithm that alternates between sequential quadratic programming updates of the admixture proportions and the allele frequencies. In addition, they use acceleration techniques [54, 58] to speed up iterations. They report considerable speedups in computing admixture proportions and population allele frequencies. However, their software, ADMIXTURE [2], only works with bi-allelic SNP data.

Our goal in this manuscript is to address lingering issues in computational efficiency, reproducibility, and reliability of genetic structure inference by developing sequential quadratic programming, and acceleration methods (*sensu* Alexander et al. [2]) for general (multiallelic) genotypic data, while accounting for the presence of missing data in our software, `MULTICLUST`. `MULTICLUST` is likelihood-based, and therefore estimates obtained from multiple models are readily amenable in a hypothesis testing or model selection framework (*e.g*., AIC, BIC, bootstrap). `MULTICLUST` estimation is also repeatable and reproducible given sufficient random initializations that altogether take a fraction of the time required by Bayesian algorithms.

## 2 Methods

### 2.1 Model

Our goal is to perform maximum likelihood estimation for the genetic mixture [50] and admixture [42] models. Here we focus on the admixture model, leaving treatment of the mixture model to the Appendix. Both models make two fundamental assumptions: Hardy-Weinberg Equilibrium (HWE) and Linkage Equilibrium (LE). HWE results when an infinite, panmictic population mates randomly without mutation, selection, migration, genetic drift, or meiotic drive. Under these conditions, each genotype is a random combination of alleles, and allele frequencies are unchanging across generations [52]. For example, letting integers represent alleles, diploid genotype {33, 2}, consisting of allele 33 and allele 2, achieves relative frequency 2*p*_33_*p*_2_, where *p*_*m*_ is the (unchanging) population allele frequency of allele *m*. HWE is reached in just one generation of mating under the above assumptions. LE posits that alleles present at different loci are also independent draws from the population. Thus, diploid, dual-locus genotype {33, 2}; {4, 17} has probability 4*p*_1,33_*p*_1,2_*p*_2,4_*p*_2,17_, for population allele frequencies now distinguishing loci 1 and 2. Under the same HWE assumptions, LE is approached at a rapid, geometric rate across generations.

If we further assume codominance of all alleles at each locus, then the ‘observed’ data are comprised of all *M* (the ploidy level) alleles of *I* individuals observed at *L* loci. Suppose these individuals derive from a population containing *K* subpopulations. Let *A*_*l*_ be the number of distinct alleles at locus *l*. For polymorphic DNA data, *A*_*l*_ = 4 and is constant across loci. For SNP data, it is typically assumed *A*_*l*_ = 2. The *m*th allele observed in individual *i* at locus *l* is denoted by *x*_*ilm*_, and all data are subsumed into the *I* × *L* × *M* array, ***X***. Because the *M* alleles observed at locus *l* are random replicates of the same process, we might restructure the data into the sufficient statistics, an *I* × *L* × {*A*_1_, …, *A*_*L*_} jagged array ***N***, where *n*_*ila*_ is the number of times the *a*th allele is observed in individual *i* at locus *l*. We caution that it is only computationally advantageous to work with the data in this form when ploidy *M* is routinely larger than the number of distinct alleles *A*_*l*_.

Subpopulations within the sampled population are distinguishable to the extent that they have different allele frequencies. Let *p*_*kla*_ be the frequency of allele *a* at locus *l* in subpopulation *k*. Clearly, 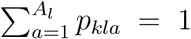 for all *k* and *l*, and 0 ≤ *p*_*kla*_ ≤ 1 for all *a*. The proportion of alleles from each of the subpopulations are governed by the admixture proportions, which may take distinct forms. Pritchard et al. [42] define the model with distinct mixing proportions ***η***_*i*_ = (*η*_*i*1_, *η*_*i*2_, …, *η*_*iK*_) for each individual *i*.

Assuming the admixture model with individual-specific mixing proportions ***η***_*i*_ and all observed alleles are independent by HWE and LE, the likelihood becomes, first in terms of data matrix ***X*** and second in terms of sufficient statistics ***N***,

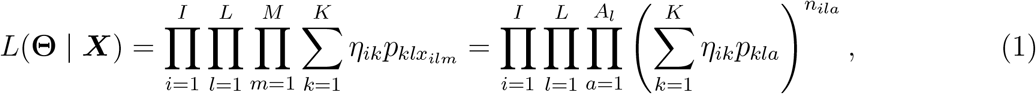

where Θ is the collection of all allele frequencies {*p*_*kla*_} and mixing proportions {*η*_*ik*_}. We allow the possibility that some alleles are not observed, or ‘missing,’ (see Appendix). Smouse et al. [50] derived the EM with missing data for the mixture model, but we believe this is the first explicit derivation for the admixture model. Though Liu et al. [29] considered missing data, they impute them from genetically similar individuals.

### 2.2 Optimization

#### 2.2.1 EM algorithm

Most authors who seek to maximize likelihood (1) (or the mixture model version (**??**)) have used the Expectation Maximization (EM) algorithm [9], with the notable exceptions of a quasi-Newton algorithm for conditional maximization assuming allele frequencies {*p*_*kla*_} known [14], a block relaxation algorithm [53], and a block relaxation algorithm with sequential quadratic programming and acceleration [2]. The EM algorithm iteratively computes the conditional expectation of the complete-data log likelihood in the E-step, followed by maximization of the expected complete-data log likelihood in the M-step until the observed log likelihood no longer increases [9]. Under the admixture model, the complete-data likelihood is the likelihood of the observed genotypic data and the unobserved population assignments of each allele, as if the latter had been observed. Our update equations (4) are formulated in terms of the allele counts ***N*** = {*n*_*ila*_}, but could be naturally adjusted to use the data matrix ***X*** = {*x*_*ilm*_}.

If we pretend to know the unobserved data, ***D*** = {*D*_*ilmk*_}, where *D*_*ilmk*_ indicates if the *m*th allele at locus *l* in individual *i* comes from the *k*th subpopulation, then the complete-data likelihood can be written as

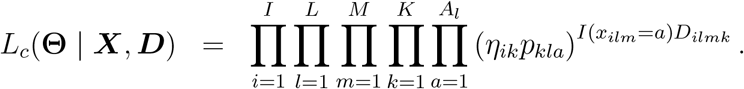

Here, the indicator function *I*(·) takes value 1 when its argument is true and is otherwise 0. During the *t*-th iteration and when *x*_*ilm*_ = *a*, we compute

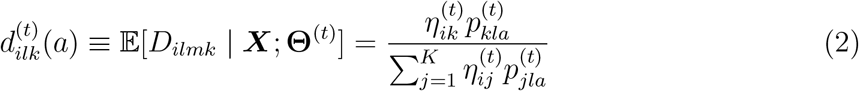

by Bayes’ rule, assuming the *t*-th parameter update **Θ**^(*t*)^. Since this expression is independent of *m*, we can rearrange sums in the expected complete-data log likelihood to get

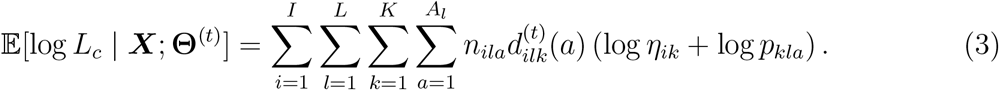

If we let 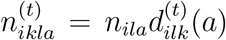 be the expected number of *a* alleles at locus *l* in individual *i* descendent from subpopulation *k* at the *t*-th iteration, then maximization of (3) yields the next (*t* + 1)-th iterates

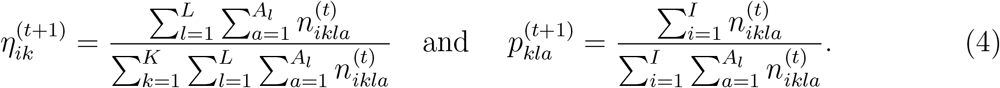

#### 2.2.2 Acceleration

The EM algorithm is an iterative procedure that can be accelerated by general numerical strategies that operate on multiple sequential iterates [54, 58]. Here, we extend the acceleration strategies already implemented in `ADMIXTURE` [2] to multi-allelic genotypes. Specifically, we implement the three SQUAREM methods [54] and the quasi-Newton acceleration methods for *q* = 1, 2, and 3 secant conditions [58].

The SQUAREM methods use update equation

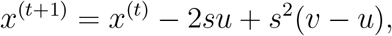

where *x*^(*t*)^ is the *t*-th iterate of the entire parameter set Θ and *u* = *F* (*x*^(*t*)^) − *x*^(*t*)^ and *v* = *F* ∘ *F* (*x*^(*t*)^) − *F* (*x*^(*t*)^) are the incremental changes in the parameter values after applying the EM algorithm, represented here as map *F* (·), twice to the last SQUAREM iterate *x*^(*t*)^. The SQUAREM methods (SE1, SE2, and SE3) differ in how they set *s* (see [54]).

The quasi-Newton method with *q* secant conditions uses update equation

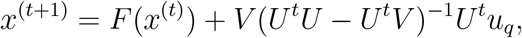

where matrices *U* = (*u*_1_, *u*_2_, …, *u*_*q*_) and *V* = (*v*_1_, *v*_2_, …, *v*_*q*_) are formed from the last *q u*_1_ = *F* (*x*^(*t−q*+1)^)−*x*^(*t−q*+1)^, …, *u*_*q*_ = *F* (*x*^(*t*)^)−*x*^(*t*)^ and *v*_1_ = *F* ∘ *F* (*x*^(*t−q*+1)^)−*F* (*x*^(*t−q*+1)^), *v*_*q*_ = *F* ∘ *F* (*x*^(*t*)^) − *F* (*x*^(*t*)^) incremental differences from two EM iterations now applied to the last *q* iterates of the quasi-Newton method.

Since accelerated updates do not necessarily follow boundary constraints, we project the results onto the appropriate simplex using the algorithm of Michelot [36]. For example, for updates to ***p***_*l*_ = {*p*_*kla*_ : 1 ≤ *k* ≤ *K*, 1 ≤ *a* ≤ *A*_*l*_}, the collectiong of allele frequencies associated with locus *l*, we project the resulting 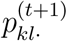 onto {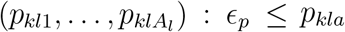 and 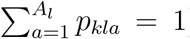} for some small *E*_*p*_ *>* 0 to avoid solutions at the boundary. For stability, we also apply the same transformation to project EM updates into the simplex. Varadhan and Roland [54] recommend adjusting the step size *s* when the SQUAREM update fails to improve the likelihood relative to the EM or BR iterate, *F* ∘ *F* (*x*^(*t*)^). We instead set *x*^(*t*+1)^ = *F* ∘ *F* (*x*^(*t*)^) when the acceleration fails for either the SQUAREM and quasi-Newton methods. Because the EM algorithm is guaranteed to increase the log likelihood, such a strategy yields an improvement each iteration even when acceleration fails.

#### 2.2.3 Initialization

Though the EM and accelerated EM are guaranteed to climb peaks in the log likelihood, not all initializations converge to the global maximum. Repeated initialization is required to find the global maximum when the likelihood surface is multimodal. We developed two schemes for initialization – random initialization (ancestry proportions sampled from a uniform distribution), and initialization based on EM (see Appendix).

### 2.3 Model Selection

#### 2.3.1 Information criteria

Choosing the number of subpopulations *K* is a model selection problem. Two information criteria, AIC = −2 log *L* + 2*p* [1] and BIC = −2 log *L* + *p* log *n* [46], are popularly used for model selection by choosing the *K* that minimizes either AIC or BIC. Here, *p* is the total number of parameters and *n* is the number of observations. For our models, we set *n* = *I* * *L* * *M* and let 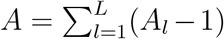, then the number of parameters is *p* = *I*(*K* − 1) + *KA* for the admixture model. Simulation studies have demonstrated that AIC tends to overestimate *K*, while BIC tends to underestimate *K* in model-based clustering [31].

#### 2.3.2 Parameteric bootstrap

If the null hypothesis is nested within the alternative hypothesis, then it might be reasonable to reject *H*_0_ in favor of *H*_1_ if the likelihood ratio *T* = −2[*l*_0_(*θ*_0_ | *X*) − *l*_1_(*θ*_1_ | *X*)] is in the right tail of the asymptotic *χ*^2^-distribution. However, the theory supporting this asymptotic distribution breaks down for mixture models [33]. Instead, bootstrapping [10] can be used to test the null hypothesis *H*_0_ : *K* = *k* vs. the alternate hypothesis *H*_1_ : *K* = *k*\* where *k* ≠ *k*\*. The parametric bootstrap generates new datasets (of multilocus genotypes) of the same size as the observed data from the fitted *H*_0_ model. The data is refit to both *H*_0_ and *H*_1_ and the likelihood ratio test statistic *T*_*b*_ is recorded for the *b*th bootstrapped dataset. The *T*_*b*_ form an empirical sampling distribution for *T* under *H*_0_, and we can estimate the *p*-value as 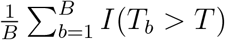. If the *p*-value is less than 0.05, then we reject *H*_0_, otherwise we take *H*_0_ as the better solution. To estimate *K*, we use *k*\* = *k* + 1 and sequentially test *H*_0_ : *K* = *k* against *H*_1_ : *K* = *k* + 1 for *k* = 1, …, *K*_max_ − 1. For each *H*_0_, we generate *B* = 100 bootstrap samples from the fitted model under *H*_0_ and compute the *p*-value given above. We conclude *K* = *k* for the smallest *k* for which *H*_0_ is not rejected. If *H*_0_ : *K* = *K*_max_ − 1 is rejected, then we conclude that *K* = *K*_max_ is statistically supported by the data. We do not adjust for multiple testing in this procedure and do not account for the possibility that a *K > k*\* is supported by the data when *H*_0_ has been accepted. Procedures for overcoming both these deficiencies are found in Maitra and Melnykov [31].

### 2.4 Simulations

We simulated ten replicate, diploid, multiallelic genotype datasets at 50 unlinked genomic loci using the coalescent simulator *msprime* v.1.0 [3] under two evolutionary demographic models with migration – (1) Island Model and (2) Stepping Stone Model – with *K* = 3, 5, and 10 subpopulations. Twenty diploid individuals(*i.e*., 40 haplotypes per subpopulation) were sampled from each subpopulation, such that the subpopulations exchanged individuals at the rate of *m* = 0.01 per generation, with unchanging population sizes of *N* = 1000. Then, *tskit* v.0.6.0 [25] was used to simulate mutations from the tree sequences generated by *msprime*, using the microsatellite mutation model, with a mutation rate of *μ* = 10^−1^. Data were then converted into the multilocus genotypic format utilized by `MULTICLUST` and `STRUCTURE` using plink 2.0 alpha [5].

Fify random initializations of `MULTICLUST` (`-n 50`) were run on each dataset (random seed specified using `-r`), varying the number of assumed clusters (`-k`) varied from 1 through *K* + 1 under the admixture model accelerated with three secant conditions (`-a -s 3`). AIC and BIC were computed from each replicate run, and used to select the best fitting model. In order to compare the performance of our acceleration algorithms against the unaccelerated EM algorithm, one such replicate dataset (under the island model, true *K* = 3) was selected and benchmarked using fifty random initializations under the admixture and mixture models with all six acceleration algorithms (`-s 1` to `-s 6`). The log likelihood and times to reach convergence were tracked under each scenario.

### 2.5 Empirical Data

We utilized a multi-allelic, multilocus genomic dataset collated from three native African populations (9 Baka, 9 Hadza, and 9 Sandawe individuals) from the study of [12] (Sarah Tishkoff, pers. comm.) to benchmark `MULTICLUST` on a large human genomic dataset. We generated a dataset containing 10,000 multi-allelic SNP’s through a pipeline that we describe elsewhere (see [48]). This dataset was then benchmarked for time under the admixture model with different acceleration schemes (`-s 1` – `-s 6`) as described above by varying the number of clusters from *K* = 1 to 3. To obtain comparable results from `STRUCTURE`, we ran this dataset of 10,000 multi-allelic SNP’s under the admixture model with a burn-in period of 10,000 and 100,000 MCMC repetitions after burn-in, assuming no linkage. The parameter *α*, which is the Dirichlet parameter that determines the degree of admixture, was inferred from the data. All other parameters were set to their default values.

To test the congruence and variances in estimates of admixture proportions and sub-population allele frequencies while utilizing `MULTICLUST` in analyzing large bi-allelic versus multi-allelic datasets, we performed replicate runs of `MULTICLUST` on the SNP dataset of [40] which analyzed haplotypic variation across 1,107 modern humans across 63 global sampling locales, genotyped at 2,810 SNPs (https://rosenberglab.stanford.edu/diversity.html). This dataset was then pruned using PPP [55] to obtain an unpruned (multi-allelic, 2,810 SNPs) and pruned (bi-allelic, 2521 SNPs) datasets, which were then analyzed with `MULTICLUST` (using the admixture model, EM accelerated with the quasi Newton algorithm with 3 secant conditions, varying K from 1 to 14, and five initializations to EM convergence – `-a -n 5 -s 3 -k k` for `k` = 1, 2, …, 14). To assess the correlation between identified clusters between the bi- and multi-allelic datasets, we computed assignment of an individual *i* to a subpopulation *k* as

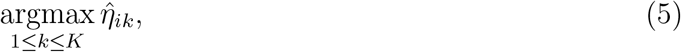

where 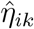 are the maximum likelihood estimated admixture proportions for individual *i*. We then computed the adjusted Rand index [43] between the two cluster assignments.

## 3 Results

### 3.1 Simulations

Under both the island model, with symmetric migration among all *K* islands, and the stepping stone model, with symmetric bidirectional migration only between adjacent islands, replicate runs of `MULTICLUST` were able to accurately recapture (Figs. 1 and 2 the true number of islands (*K* = 3, 5, 10), with decreasing accuracy and increased variance in AIC estimates with increased number of islands (*K*).

**Figure 1:**
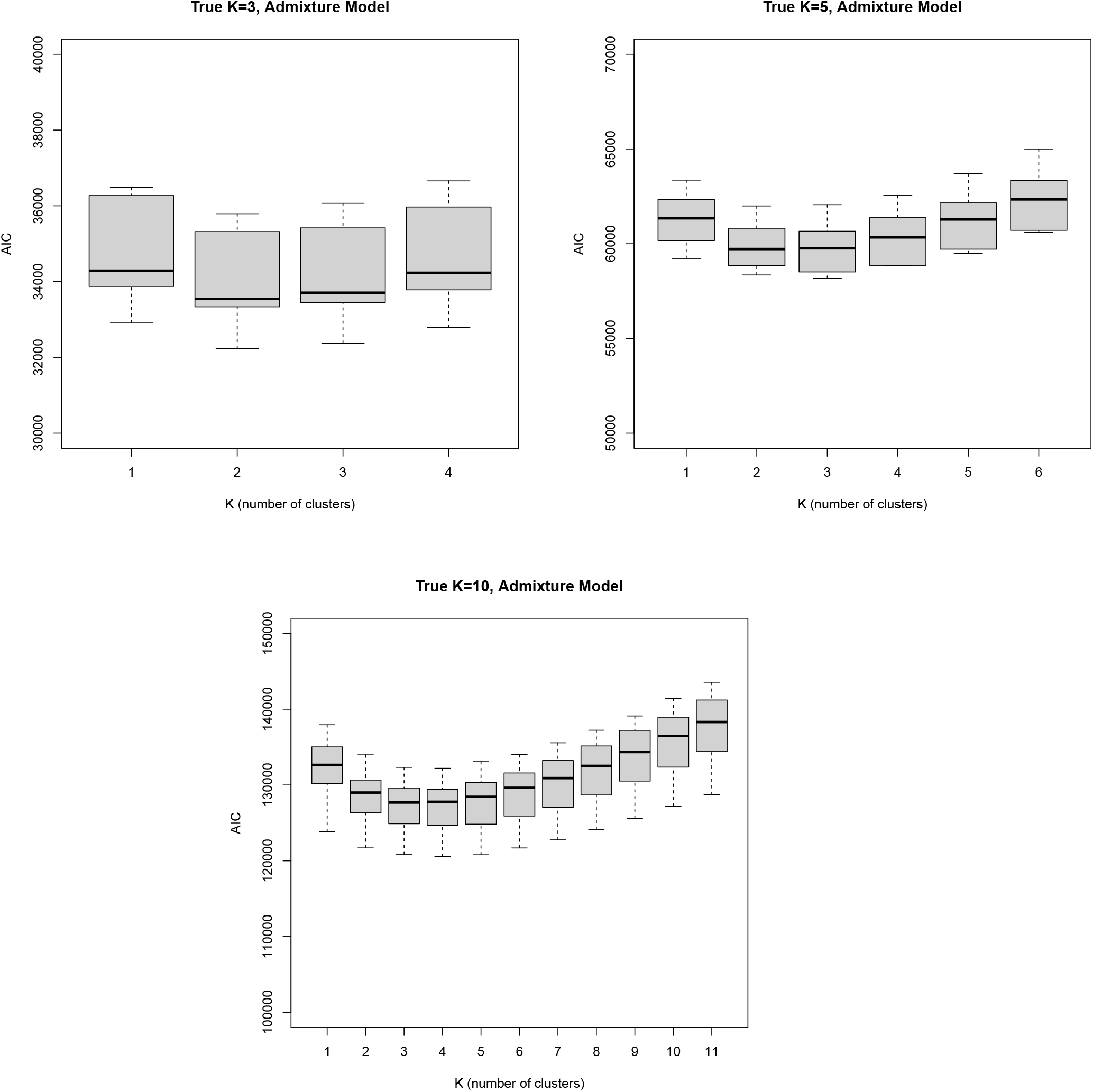
Estimates of AIC from 20 replicate runs of `MULTICLUST` under the admixture model by varying *K*. Multi-allelic genomic data at 50 loci were simulated under the Island Model with migration (assuming true *K* = 3, 5, 10).

**Figure 2:**
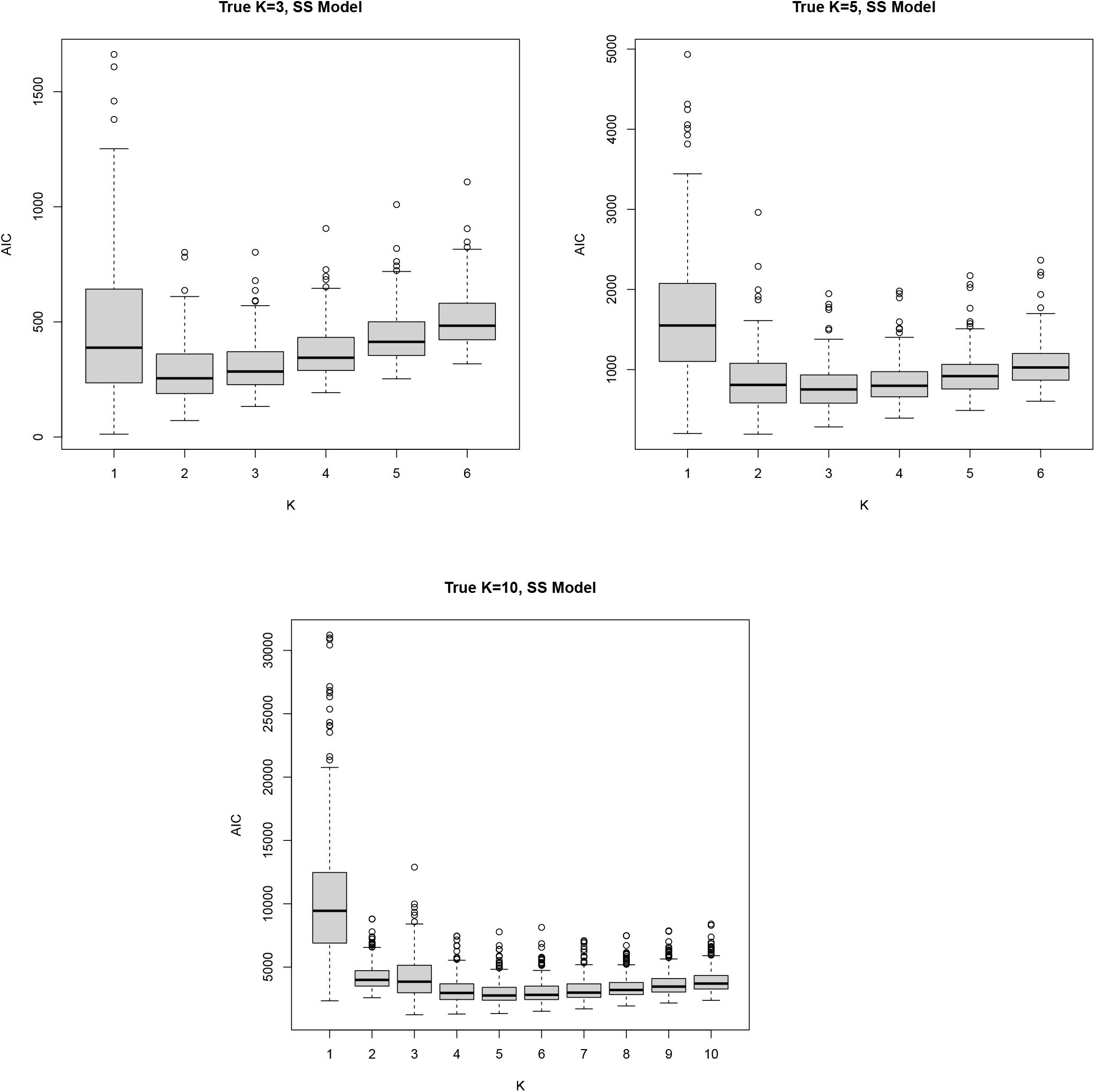
Estimates of AIC from 20 replicate runs of `MULTICLUST` under the admixture model by varying *K*. Multi-allelic genomic data at 50 loci were simulated under the Stepping Stone (SS) Model with migration (assuming true *K* = 3, 5, 10).

Across replicate *K* = 3 island model with migration runs (with varying initializations), the accelerated runs outperformed the unaccelerated EM algorithm runs in computation times (see Fig. 3). The accelerated EM algorithm offered anywhere from 3 − 30% improvement in computational time (compared to the unaccelerated EM algorithm) under the admixture model. The mixture model runs had comparable run times across accelerated and unaccelerated replicate runs.

**Figure 3:**
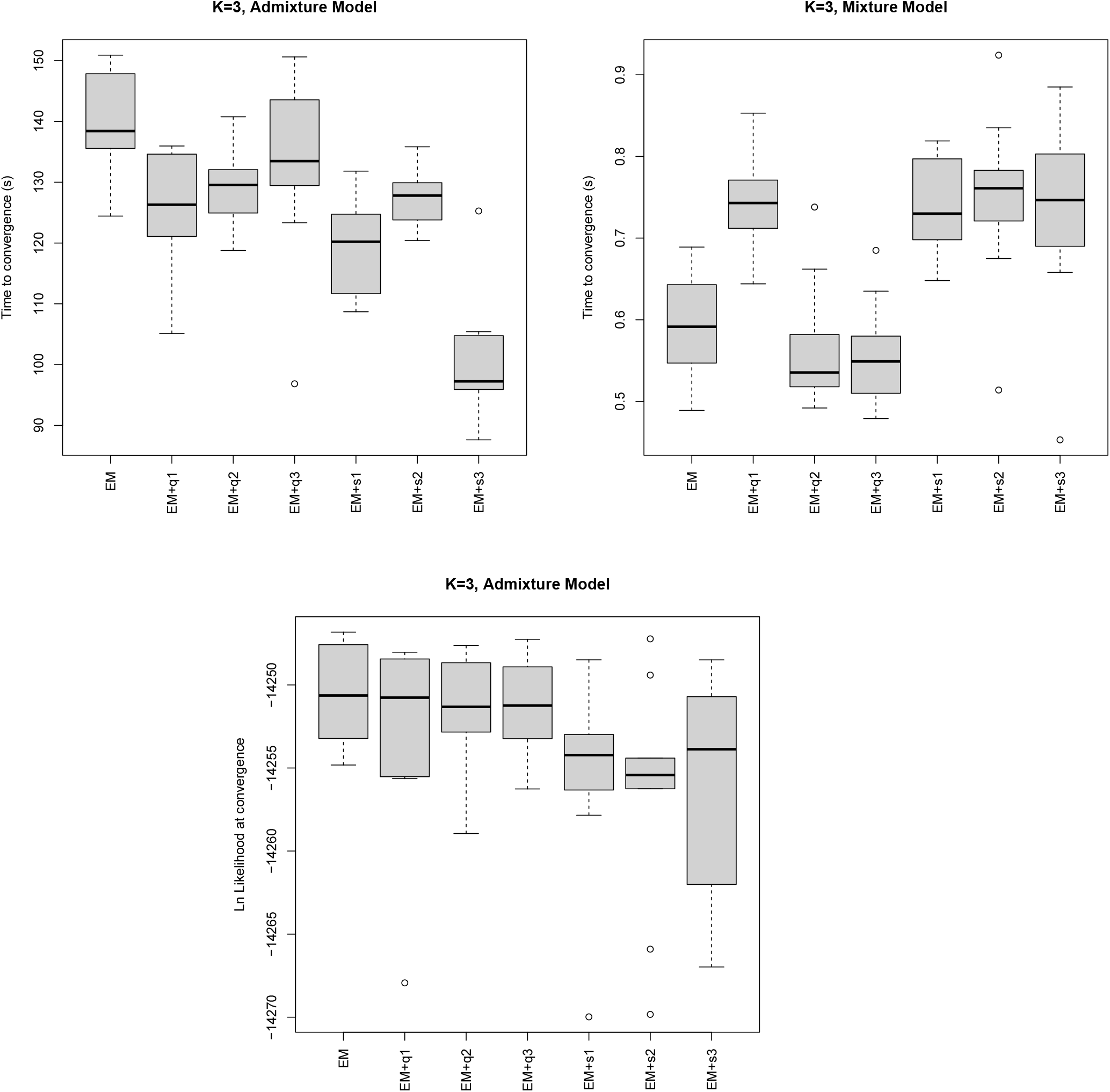
(Top) Time to convergence and (Bottom) ln likelihood at convergence from replicate runs of MULTICLUST under the admixture and mixture models, assuming *K* = 3. Benchmarking was performed across all the algorithms implemented in MULTICLUST - EM = unaccelerated EM algorithm, EM + q1/q2/q3 = EM algorithm, accelerated via SQUAREM versions 1-3, and EM + s1/s2/s3 = EM algorithm, accelerated via the Quasi Newton algorithm with 1-3 secant conditions respectively. Multi-allelic genomic data were simulated for these analyses under the island model with migration, with a true *K* = 3.

Absolute difference in logarithmic likelihood between alternate runs of unaccelerated and accelerated EM runs ranged from indicated convergence to the same maxima by all algorithms alike (admixture model: mean ln likelihood = −14253.528, std. dev. = 5.561, mixture model: mean ln likelihood = −14691.866, std. dev. = 48.199).

### 3.2 Empirical Data

Both the accelerated and unaccelerated EM runs from `MULTICLUST` on the African dataset (true *K* = 3) accurately estimated population structure at *K* = 3 using AIC. All algorithms also converged within ~ 1 minute of run time on a single node of a 16-node Dual-10 Xeon CPU, E5-2630v4 2.20GHz, 12.8GB RAM per core machine using `MULTICLUST`. Acceleration using three secant conditions (`-q3`) offered the greatest decrease (41%) in run time. Comparable runs of `STRUCTURE` failed to converge past over 15 minutes of run time on the same node (Table 1).

**Table 1:** Maximum log likelihood at convergence and time to convergence (in minutes) estimated from a dataset containing 10,000 multi-allelic SNPs derived from three native African populations from the 1000 Genomes Project at *K* = 2, 3 – estimates from `MULTICLUST` (EM, acceleration schemes) are presented in comparison with long `STRUCTURE` [11, 42] runs that were terminated after 15 minutes/20 minutes respectively.

| Algorithm | $K = 2$ | | $K = 3$ | |
| --- | --- | --- | --- | --- |
|  | Likelihood | Time | Likelihood | Time |
| STRUCTURE | −93 185.60 | 15 h 22 m | −93 402.00 | 20 h 47 m |
| EM only | −84 599.76 | 1 m 9 s | −79 995.75 | 1 m 22 s |
| EM + s1 | −84 599.75 | 42 s | −79 995.65 | 34 s |
| EM + s2 | −84 599.84 | 50 s | −79 996.11 | 42 s |
| EM + s3 | −84 599.84 | 33 s | −79 996.30 | 32 s |
| EM + q1 | −84 600.51 | 45 s | −79 996.62 | 33 s |
| EM + q2 | −84 599.91 | 47 s | −79 995.50 | 1 m 2 s |
| EM + q3 | −84 600.09 | 46 s | −80 003.26 | 34 s |

Replicate runs of `MULTICLUST` under the admixture model with acceleration (`-q3`) by varying *K* from 1–14 on the larger bi-allelic versus multi-allelic HGDP dataset of [40] (Table 2) showed decreasing concordance in clustering with increasing *K*. Lower Rand indices are however obtained between bi-allelic and multi-allelic datasets with greater *K*, which are most likely due to initialization issues with convex optimization. This issue will be discussed further below. Estimated admixture proportions using the multi-allelic dataset are shown in Fig. 4, which is largely consistent with expected geographical structure and admixture across modern human populations.

**Figure 4:**
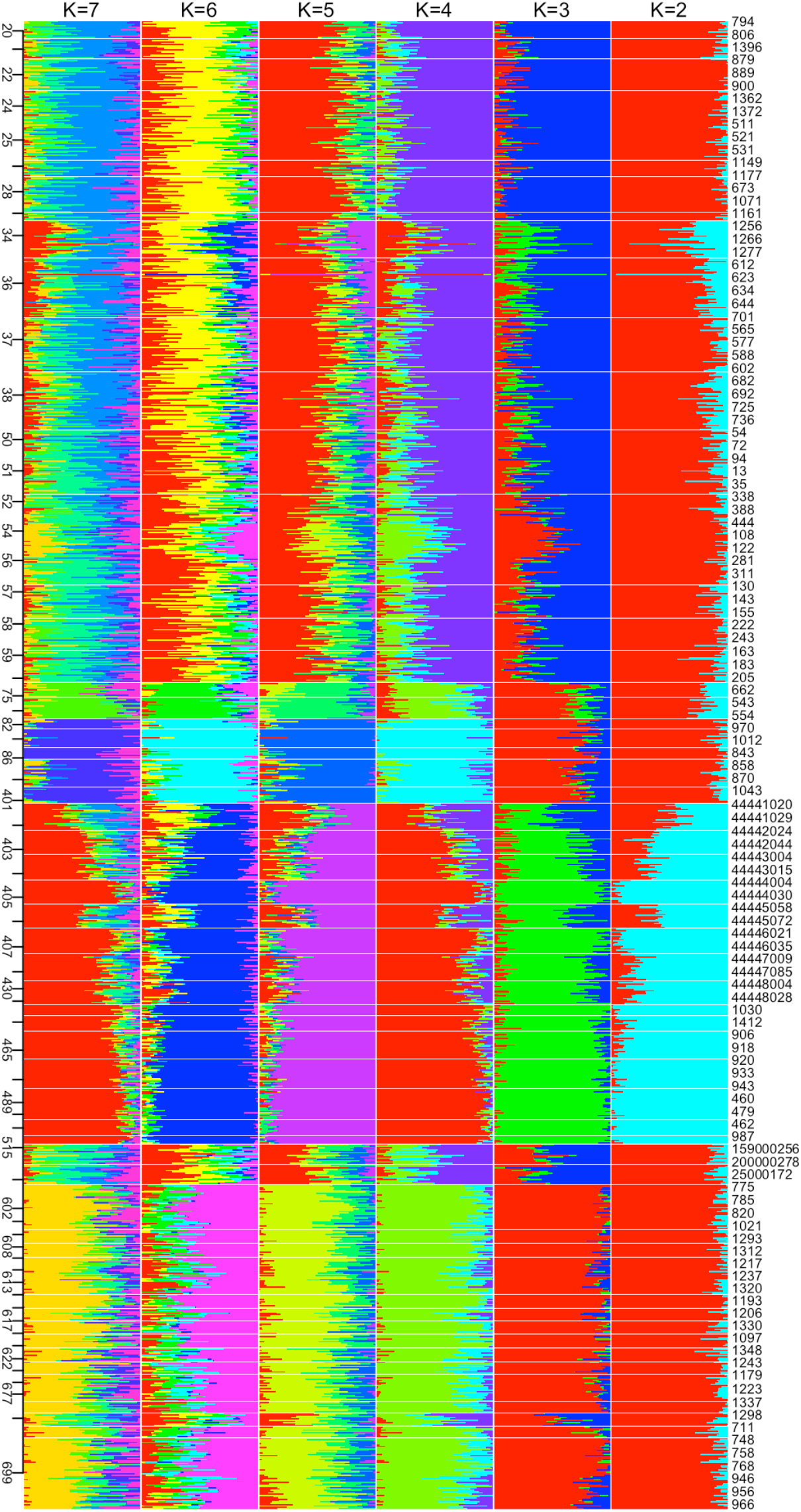
Stacked barplots of admixture proportions of global human populations, estimated via replicate runs of MULTICLUST assuming *K* = 2, …, 7 from the HGDP dataset of Pemberton et al., 2013. Population ID’s are reported on the left, while individual ID’s are reported on the right, and obtained from https://rosenberglab.stanford.edu/diversity.html. Global population structure of modern humans estimated using MULTICLUST are concordant with geographical structure, as reported in previous studies.

**Table 2:** Estimates of AIC and BIC from 50 replicate runs of `MULTICLUST` on the HGDP dataset of Pemberton et al. [41], comprising 2,810 single-nucleotide polymorphisms in 1,107 individuals from 63 populations. Data were pruned to only include biallelic loci, and compared against the full dataset with multiallelic loci. Also reported are the Adjusted Rand Indix (ARI) between the clusterings (computed via the maximum ancestry proportion across *K* clusters) from the bi-allelic versus multi-allelic datasets. Increased number of assumed clusters (*K*) results in greater discordance amongst assigned clusters.

| *K* | Biallelic |  | Multiallelic |  | ARI |
| --- | --- | --- | --- | --- | --- |
|  | AIC | BIC | AIC | BIC |  |
| 1 | 5 758 103.67 | 5 770 732.39 | 6 139 359.85 | 6 154 884.01 | 1.00 |
| 2 | 5 537 488.93 | 5 568 291.79 | 5 904 590.18 | 5 941 183.92 | 1.00 |
| 3 | 5 377 063.15 | 5 426 040.14 | 5 732 676.60 | 5 790 339.90 | 0.97 |
| 4 | 5 318 800.51 | 5 385 951.64 | 5 671 406.96 | 5 750 139.84 | 0.97 |
| 5 | 5 283 651.01 | 5 368 976.27 | 5 634 705.40 | 5 734 507.85 | 0.97 |
| 6 | 5 255 492.43 | 5 358 991.83 | 5 604 667.21 | 5 725 539.24 | 0.82 |
| 7 | 5 233 921.20 | 5 355 594.73 | 5 583 270.66 | 5 725 212.26 | 0.82 |
| 8 | 5 213 973.46 | 5 353 821.13 | 5 564 560.35 | 5 727 571.53 | 0.63 |
| 9 | 5 198 327.36 | 5 356 349.16 | 5 548 423.88 | 5 732 504.63 | 0.65 |
| 10 | 5 183 896.14 | 5 360 092.08 | 5 535 834.73 | 5 740 985.06 | 0.56 |
| 11 | 5 170 213.80 | 5 364 583.88 | 5 520 466.89 | 5 746 686.79 | 0.65 |
| 12 | 5 159 103.16 | 5 371 647.37 | 5 513 531.30 | 5 760 820.77 | 0.58 |
| 13 | 5 151 952.93 | 5 382 671.27 | 5 502 198.02 | 5 770 557.07 | 0.54 |
| 14 | 5 141 102.06 | 5 389 994.54 | 5 492 371.87 | 5 781 800.49 | 0.59 |

## 4 Discussion

Model-based genetic clustering methods of interest are simple models that assume that sampled individuals originate from *K* subpopulations, each characterized by distinct allele frequencies. In the *mixture* model, each individual derives from just one of the *K* possible subpopulations, so all observed alleles of that individual are independently drawn from the same source. In the *admixture* model, each allele within an individual is sampled independently from any of the *K* subpopulations. In the latter model, an individual is characterized by admixture proportions that determine the fraction of alleles derived from each subpopulation. Neither model is particularly realistic; they are useful approximations to much more complex realities. The mixture model technically describes a sample taken from *K* cryptic subpopulations that are not interbreeding, and was originally used for attributing fish catch to breeding grounds [50]. The admixture model is appropriate for populations formed from the merger of *K* subpopulations with sufficient subsequent interbreeding to eliminate admixture linkage [11]. The general admixture model allows each individual to be a distinct “admixture” of source populations, which exactly matches no physical process, but perhaps reasonably mimics the individual-specific admixtures that may result from varying spatial proximity to source populations or restricted interbreeding patterns among ancestral populations.

Several methods have been developed to infer genetic population mixtures using multilocus genotype data [2, 42, 53]. These methods can be slow, especially when used to estimate the number of subpopulations (*K*) for larger *K*, or for larger datasets, which often requires repeated runs. Perhaps as a consequence of insufficient replication, these methods often make unreliable inference on the ‘true’ value of *K*. Three major issues arise in utilizing these tools - (a) Inferring the true value of *K* ancestral subpopulations, (b) Inconsistencies in inferred true values of *K* over multiple iterations (and initializations) of these tools, and (c) Speed of computations, leading to limitations on testing greater values of *K*, and for larger datasets (with more individuals or loci or alleles per locus). `MULTICLUST` extends the acceleration strategies of Alexander et al. [2] to multiallele traits and outperforms `STRUCTURE` [42] for estimation of *K* with respect to speed of computation.

Over all the tests we performed with simulated and empirical data, we obtained comparable or better (more congruence with ‘true’ structure) results.

Despite the speed-ups offered by the accelerated EM algorithms, we have demonstrated both methods with simulated and empirical data to be at least as accurate as each other.

Inference of both model parameters and *K* is stochastic for all methods, and as a result multiple runs of the software, even with the same settings, may produce different estimates. It is well known that improper run settings can lead to lack of reproducibility in Bayesian inference by MCMC. Traditionally, estimation process is allowed to proceed for a certain ‘burn-in’ period, all estimates of which are discarded prior to the actual MCMC iterations that are retained and corresponding parameter estimates are obtained from. There has been considerable debate on how long these burn-in periods and how many MCMC repetitions are needed for inference (see Gilbert et al. [18], Latch et al. [26], Tutorial for `STRUCTURE`, BAPS, etc), with the general consensus that the longer, the better. Meanwhile maximum likelihood methods find local maxima, and may produce different results across initializations, which requires multiple initializations to be performed to obtain better estimates. We used an arbitrary number (50) initializations across all our tests of the accuracy of both the EM and BR algorithms. In light of the importance of repetition, the speed of inference becomes critical. Other initialization strategies, such as K-means/medoids would perhaps lead to faster estimates using both algorithms, but is yet to be explored. We have also shown that the accelerated EM algorithms require fewer initializations to converge at the same maxima as the EM algorithm.

One key utility of `MULTICLUST` is that we often also obtained these results in a fraction of the time required to obtain convergence from `STRUCTURE` – a major hurdle for ‘problematic’ datasets, where inference of *K* is difficult, either owing to great degrees of admixture or inflated parameter sets due to excessive heterozygosity and/or a large number of observed alleles at multiple loci. For instance, inference of the true number of subpopulations is challenging under evolutionary scenarios such as the Stepping Stone Model (a discrete approximation of the Isolation By Distance). In these scenarios, even though adjacent populations are more likely to be structured together into the same ancestral subpopulation, inference of true *K* becomes muddled owing to the distribution of allele frequencies over a continuum of (sub)populations. On the other hand, inference of true *K* becomes easier in populations that are significantly differentiated, with several alleles being fixed to localized subpopulations (eg. the Hierarchical Island Model).

But regardless of speed and efficiency of computation, the issues of difficulty in inferring *K* persist with `STRUCTURE` and `MULTICLUST` for the same reasons of model assumptions on equilibria and sampling distribution of allele counts. A more exhaustive sequential bootstrap methods to assess significance of clustering, as suggested by Maitra and Melnykov [31] would likely be good solutions for larger, more difficult datasets, where conflicting AIC and BIC, or if the regular non-parametric bootstrap do not yield definitive solutions. Equilibrium assumptions are also yet to be relaxed for model-based clustering methods to identify subpopulation structure. One assumption, that of linkage equilibrium (LE) was relaxed and implemented into the admixture model by Falush et al. [11], and subsequent versions of `STRUCTURE` [42]. Of future research would be to incorporate chromosomal linkage into the likelihood framework, described in this manuscript. `MULTICLUST` also assumes that alleles are sampled from a randomly mating population in HWE, which follows a simple multinomial distribution. Natural populations need not necessarily be randomly mating, which could give rise to exceptions to the multinomial sampling of allele counts. This issue is addressed in `STRUCTURE` [42], where allele counts are sampled according to probabilities drawn from a Dirichlet distribution instead, with a parameter, *α*. Of potential interest is a Dirichlet extension of `MULTICLUST`, to account for exceptions to model assumptions on random mating.

Additionally, we caution that the admixture model assumptions are rarely satisfied by real biological populations. It is important to note that the LE assumption implies that the interbreeding accompanying a population admixture has continued for enough generations to eliminate the linkage disequilibrium (LD) induced by mixing populations, so-called admixture linkage [11]. Perhaps the most biologically realistic of the admixture model variants (`STRUCTURE`’s admixture, *constrained admixture*, and *overparameterized admixture*), is the *constrained admixture*, which arises after founding a population from *K* sources (subpopulations) mixed in proportions ***η*** and then ‘breeding’ it according to the HWE assumptions until LE is reestablished. The other models are misspecifications of more complex realities and hence only approximate the true structure in the data. Probably the most serious of these approximations is LE, especially given dense genomic datasets made feasible via third generation sequencing technologies. Falush et al. [11] was the first to consider relaxation of this assumption, but there have been many subsequent methods, mostly focusing on situations relevant to human populations [19, 30].

## 5 Acknowledgments

This work was supported by a James Cornette Fellowship from the Bioinformatics and Computational Biology Program at Iowa State University, NSF CAREER 2147812 and NIH 1R15GM143700-01 to AS. All analyses were performed on the Temple University HPC (Owlsnest2), which was supported in part by the NSF major research instrumentation grant number CNS-09-58854, and on the San Diego State University HPC (mesxuuyan), which was supported by startup funds to AS and NSF-ABI: 1564659 to AS. We thank members of the Janzen lab (at ISU), and the Hey lab (at Temple University) for comments on early versions of this manuscript. We also thank David Alexander, John Novembre, and Hua Zhou for suggestions on the quadratic programming updates. We thank Sarah Tishkoff for access to the native African genomic data.

## 6 Data Availability

`MULTICLUST` is available on GitHub and can be used under GPL-3.0 license at https://github.com/arunsethuraman/multiclust. All simulated datasets are available at https://github.com/ClassicalKooz/Multiclust_Simulations.

