## Appendix for "MULTICLUST – Fast Multinomial Clustering of multiallelic genotypes to infer genetic population structure"

July 29, 2025

### 1 APPENDIX

#### 1.1 EM Algorithm

In general, the Expectation-Maximization (EM) algorithm works by iteratively computing the conditional expectation of the complete-data log likelihood in the E-step, followed by maximization of expected complete-data log likelihood in the M-step until the observed log likelihood no longer increases [2]. The complete-data likelihood is the likelihood of the observed data and the missing data. The missing data are the population assignments of each individual, in the mixture model, or allele, in the admixture model. The missing data may also include literal missing allele observations when the data are incompletely observed. Our derivations are formulated in terms of the allele counts $\boldsymbol{N} = \{n_{ila}\}$, but could be naturally adjusted to use the data matrix $\boldsymbol{X} = \{x_{ilm}\}$.

##### 1.1.1 Mixture Model

If we pretend to know $V_{ik}$, which indicates if the individual $i$ comes from the $k$th subpopulation, then the complete-data likelihood is

$$L_c(\mathbf{\Theta}, \boldsymbol{V} \mid \boldsymbol{X}) \;=\; \prod_{i=1}^{I}\prod_{k=1}^{K}\left[\eta_k \prod_{l=1}^{L}\prod_{a=1}^{A_l} p_{kla}^{n_{ila}}\right]^{V_{ik}}. \tag{1}$$

For iteration $t+1$, the E-step needs the conditional expectation of matrix $\boldsymbol{V} = [V_{ik}]_{I\times K}$, given the current parameter estimate $\mathbf{\Theta}^{(t)}$ and the observed data $\boldsymbol{X}$. Let $v_{ik}^{(t)} = \mathbb{E}[V_{ik} \mid \mathbf{\Theta}^{(t)}, \boldsymbol{X}]$. Then,

$$v_{ik}^{(t)} = \frac{\eta_k^{(t)} \prod_{l=1}^{L}\prod_{a=1}^{A_l}(p_{kla}^{(t)})^{n_{ila}}}{\sum_{j=1}^{K}\eta_j^{(t)}\prod_{l=1}^{L}\prod_{a=1}^{A_l}(p_{jla}^{(t)})^{n_{ila}}}. \tag{2}$$

The M-step maximizes the conditional expectation of the complete-data log likelihood, producing

$$\eta_k^{(t+1)} = \frac{\sum_{i=1}^{I} v_{ik}^{(t)}}{I} \quad \text{and} \quad p_{kla}^{(t+1)} = \frac{\sum_{i=1}^{I} v_{ik}^{(t)} n_{ila}}{M\sum_{i=1}^{I} v_{ik}^{(t)}}.$$

The derivations follow from the standard EM algorithm for mixture models [3].

##### 1.1.2 Admixture Model

If we pretend to know $D_{ilmk}$, which indicates if the $m$th allele at locus $l$ in individual $i$ comes from the $k$th subpopulation, then the complete-data likelihood can be written as

$$L_c(\mathbf{\Theta}, \boldsymbol{D} \mid \boldsymbol{X}) \;=\; \prod_{i=1}^{I}\prod_{l=1}^{L}\prod_{m=1}^{M}\prod_{k=1}^{K}\prod_{a=1}^{A_l}(\eta_{ik}p_{kla})^{I(x_{ilm}=a)D_{ilmk}}.$$

Here, we introduce indicator function $I(\cdot)$, which takes value 1 when its argument is true and is otherwise 0. When $x_{ilm} = a$, we compute

$$d_{ilk}^{(t)}(a) \equiv \mathbb{E}[D_{ilmk} \mid \mathbf{\Theta}^{(t)}, \boldsymbol{X}] = \frac{\eta_{ik}^{(t)} p_{kla}^{(t)}}{\sum_{j=1}^{K}\eta_{ij}^{(t)} p_{jla}^{(t)}} \tag{3}$$

by Bayes' rule. Since this expression is independent of $m$, we can rearrange sums in the expected complete-data log likelihood to get

$$\mathbb{E}[\log L_c \mid \boldsymbol{\Theta}^{(t)}, \boldsymbol{X}] = \sum_{i=1}^{I} \sum_{l=1}^{L} \sum_{k=1}^{K} \sum_{a=1}^{A_l} n_{ila} d_{ilk}^{(t)}(a) \left( \log \eta_{ik} + \log p_{kla} \right). \tag{4}$$

If we let $n_{ikla}^{(t)} = n_{ila} d_{ilk}^{(t)}(a)$ be the expected number of $a$ alleles at locus $l$ in individual $i$ descendent from group $k$ at the $t$th iteration, then maximization of (4) yields

$$\eta_{ik}^{(t+1)} = \frac{\sum_{l=1}^{L} \sum_{a=1}^{A_l} n_{ikla}^{(t)}}{\sum_{k=1}^{K} \sum_{l=1}^{L} \sum_{a=1}^{A_l} n_{ikla}^{(t)}} \quad \text{and} \quad p_{kla}^{(t+1)} = \frac{\sum_{i=1}^{I} n_{ikla}^{(t)}}{\sum_{i=1}^{I} \sum_{a=1}^{A_l} n_{ikla}^{(t)}}. \tag{5}$$

#### 1.2 Missing Values

Genetic data are replete with missing information. As Smouse et al. [5] does for the mixture model and Tang et al. [6] does for the admixture model, we assume the missing data is missing completely at random and is ignorable in Rubin's sense [4]. The genetic data can be split into observed and missing parts, $\boldsymbol{X} = \{\boldsymbol{X}^{\text{obs}}, \boldsymbol{X}^{\text{mis}}\}$. The likelihood function becomes

$$L(\boldsymbol{\Theta} \mid \boldsymbol{X}^{\text{obs}}) = \sum_{\boldsymbol{X}^{\text{mis}}} L(\boldsymbol{\Theta}, \boldsymbol{X}^{\text{mis}} \mid \boldsymbol{X}^{\text{obs}}).$$

In the E-step of EM algorithm, we need to calculate the expectation over the missing alleles as well as the unknown population assignments.

##### 1.2.1 Mixture Model with Missing Data

We will typically signal missing data by encoding missing $x_{ilm}$ with a special value, such as $-9$, 0 or '–'. Then, the observed data likelihood is unchanged from (**??**), if we understand $p_{kl0}$, for example, to be 1. The complete-data likelihood includes random variables $V_{ik}$ and $I(x_{ilm} = a)$ when $x_{ilm}$ is missing, so the expected complete-data log likelihood needed for the E-step is

$$E\left[l_c(\boldsymbol{\Theta} \mid \boldsymbol{V}, \boldsymbol{X}) \mid \boldsymbol{X}^{\text{obs}}, \boldsymbol{\Theta}^{(t)}\right] = \sum_{i=1}^{I}\sum_{k=1}^{K} v_{ik}^{(t)} \left\{ \log \eta_k + \sum_{l=1}^{L}\sum_{a=1}^{A_l} \left[ n_{ila} + n_{il0} p_{kla}^{(t)} \right] \log p_{kla} \right\},$$

where $v_{ik}^{(t)}$ is as in the case with no missing data, and $n_{il0}$ is the number of missing alleles in individual $i$ at locus $l$. We have used the independence of alleles within an individual, so conditional probability $P(x_{ilm} = a \mid V_{ik} = 1, \boldsymbol{X}^{\text{obs}}, \boldsymbol{\Theta}^{(t)})$ is $p_{kla}^{(t)}$. The M-step yields the very same update for $\eta_k^{(t+1)}$, but now

$$p_{kla}^{(t+1)} = \frac{\sum_{i=1}^{I} v_{ik}^{(t)} (n_{ila} + n_{il0} p_{kla}^{(t)})}{M \sum_{i=1}^{I} v_{ik}^{(t)}}.$$

##### 1.2.2 Admixture Model with Missing Data

The expected complete-data log likelihood for the admixture model is

$$\sum_{i=1}^{I}\sum_{l=1}^{L}\sum_{k=1}^{K}\sum_{a=1}^{A_l} \left( n_{ila} d_{ilk}^{(t)}(a) + n_{il0} \eta_{ik}^{(t)} p_{kla}^{(t)} \right) (\log \eta_{ik} + \log p_{kla}).$$

This equation is derived by recognizing that $\mathbb{E}\left[D_{ilmk} I(x_{ilm} = a) \mid \boldsymbol{X}^{\text{obs}}, \boldsymbol{\Theta}^{(t)}\right]$ is the previously computed $d_{ilk}^{(t)}(a)$ of Eq. (3) when $x_{ilm} = a$ is observed and $\eta_{ik}^{(t)} p_{kla}^{(t)}$ when $x_{ilm}$ is missing. The M-step equations (5) are valid if we define $n_{ilka}^{(t)} = n_{ila} d_{ilk}^{(t)}(a) + n_{il0} \eta_{ik}^{(t)} p_{kla}^{(t)}$.

### 2 References

**Algorithm 1:** Acceleration algorithm proposed by Varadhan and Roland [7] and Zhou et al. [8], also implemented in Alexander et al. [1].

**begin**
  **repeat**
    Run EM or BR twice to get $\eta_{ik}$ and $p_{kla}$, and save estimates for all alleles and individuals.
    **for** *;* **do**
    **repeat**
      Calculate log likelihood $L_1$
      **if** *Admixture Model* **then**
        Set $X_0 = \{\eta_{i1}, \eta_{i2}, \ldots, \eta_{ik}, p_{kl1}, p_{kl2}, \ldots, p_{klm}\}$ from previous iteration
        Set $X_1 = \{\eta_{i1}, \eta_{i2}, \ldots, \eta_{ik}, p_{kl1}, p_{kl2}, \ldots, p_{klm}\}$ from iteration 1 of EM/BR
        Set $X_2 = \{\eta_{i1}, \eta_{i2}, \ldots, \eta_{ik}, p_{kl1}, p_{kl2}, \ldots, p_{klm}\}$ from iteration 2 of EM/BR
      **else if** *Mixture Model* **then**
        Set $X_0 = \{\eta_1, \eta_2, \ldots, \eta_k, p_{kl1}, p_{kl2}, \ldots, p_{klm}\}$ from previous iteration
        Set $X_1 = \{\eta_1, \eta_2, \ldots, \eta_k, p_{kl1}, p_{kl2}, \ldots, p_{klm}\}$ from iteration 1 of EM/BR
        Set $X_2 = \{\eta_1, \eta_2, \ldots, \eta_k, p_{kl1}, p_{kl2}, \ldots, p_{klm}\}$ from iteration 2 of EM/BR
      Calculate $U = X_1 - X_0$ and $V = X_2 - X_1$
      Calculate step size $s$
      Compute $X_{n+1}$
      Calculate log likelihood $L_2$
    **until** $L_1 > L_2$
  **until** $L_2 - L_1 < 10^{-4}$
